# RIBOSS detects novel translational events by combining long- and short-read transcriptome and translatome profiling

**DOI:** 10.1101/2024.11.07.622529

**Authors:** Chun Shen Lim, Chris M. Brown

## Abstract

Ribosome profiling is a high-throughput sequencing technique that captures the positions of translating ribosomes on RNAs. Recent advancements in ribosome profiling include achieving highly-phased ribosome footprints for plant translatomes and more recently for bacterial translatomes. This substantially increases the specificity of detecting open reading frames (ORFs) that can be translated, such as small ORFs located upstream and downstream of the annotated ORFs. However, most genomes (e.g. bacterial genomes) lack the annotations for the transcription start and termination sites. This hinders the systematic discovery of novel ORFs in the ‘untranslated’ regions in ribosome profiling data. Here we develop a new computational pipeline called RIBOSS. We use RIBOSS to leverage long-read and short-read data for *de novo* transcriptome assembly, and highly-phased ribosome profiling data for detecting novel translational events in the newly assembled transcriptome. We demonstrate the capability of RIBOSS using recently published metatranscriptome and translatome data for *Salmonella enterica* serovar Typhimurium. The RIBOSS Python modules are versatile and can be used to analyse prokaryotic or eukaryotic data. In sum, RIBOSS is the first computational pipeline to integrate long- and short-read sequencing technologies to investigate translation. RIBOSS is freely available at https://github.com/lcscs12345/riboss.

## INTRODUCTION

Ribosome profiling is a powerful technique for detecting the positions of ribosomes on open reading frames (ORFs) (*1, 2*). The standard ribosome profiling technique employs RNase to digest the ribosome-bound RNAs and sucrose gradients to isolate single ribosomes with mRNA fragments protected. Ribosome profiling has been improved over the years and achieved ‘super-resolution’, notably in plant translatomes (*3, 4*). Recently, the human ORFs discovered through ribosome profiling have been curated and released as a catalogue for reference gene annotation projects (*5*). This community-led effort has been endorsed by the experts in the HUPO/HPP Executive Committee, Human Technopole, and RIKEN. These contributions have significantly improved our understanding of cellular functions.

However, gene annotation for bacteria still lags behind. Highly phased ribosome profiling data for bacteria has only recently become available (*6*). This opens the possibility of detecting novel translational events in bacteria with high confidence. However, the transcription start and termination sites and the intercistronic regions in many important pathogenic bacteria have yet to be determined and identified. These regions can also harbour unusual translational events (*7, 8*).

To overcome this problem, we have developed a new computational pipeline called RIBOSS, which combines long- and short-read RNA sequencing data for *de novo* transcriptome assembly, and ribosome profiling data to identify novel translational events in the assembled transcriptome. RIBOSS uses state-of-the-art software for long- and short-read alignment, transcriptome assembly, and transcript and ribosome footprint quantification. RIBOSS includes six modules tailored for ribosome footprint analysis, and statistical analysis for comparing the translation of novel ORFs with nearby annotated ORFs.

## RESULTS AND DISCUSSION

### RIBOSS combines long- and short-read data for transcriptome assembly with highly-phased ribosome profiling data

We support the notion that the completeness of gene annotation is critical for understanding biological functions ( *5* ). However, most current gene annotation pipelines, especially for prokaryotes, annotate only the protein-coding regions and omit the transcription start and termination sites and the intercistronic regions (e.g. NCBI PGAP and Prokka) ( *9* – *11* ). This hampers the discovery of novel translational events beyond annotated ORFs.

Therefore, we have developed RIBOSS to harness long-read sequencing data for *de novo* transcriptome assembly, and high-quality ribosome profiling data for detecting novel ORFs in the newly assembled transcripts (Fig 1). To demonstrate how RIBOSS works, we have analysed the transcriptome and translatome data for *Salmonella enterica* serovar Typhimurium ( *S* . Typhimurium).

**Fig 1.**
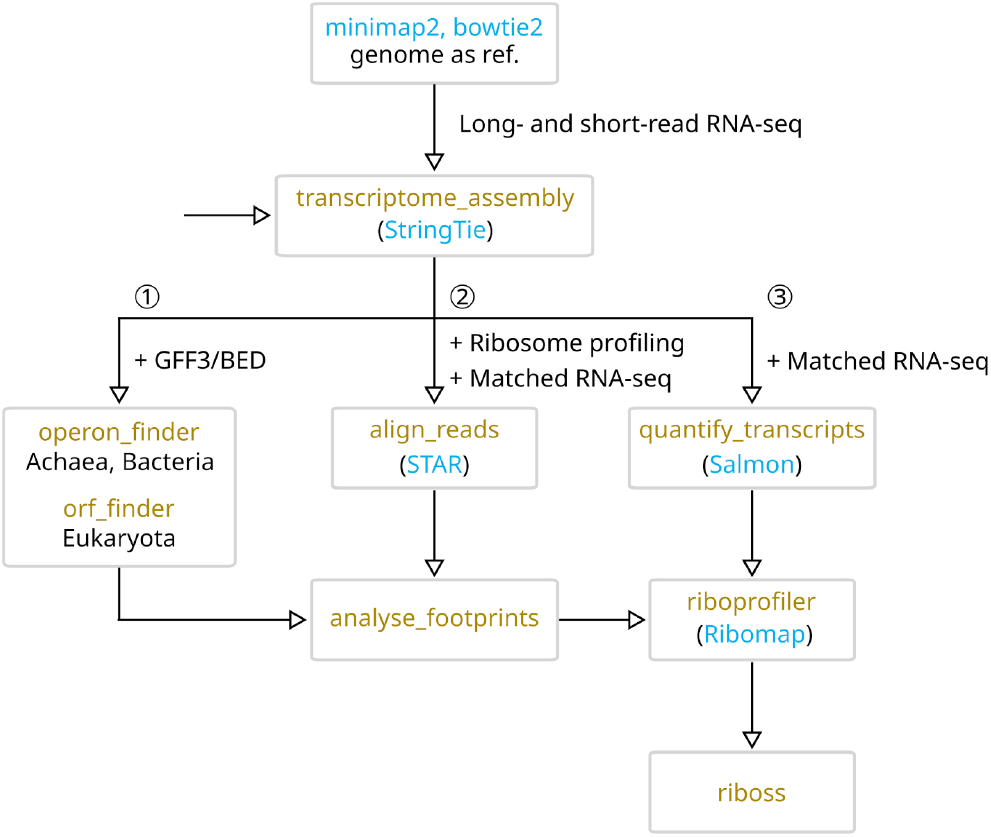
Analysis pipeline for long-short read transcriptome and translatome data using RIBOSS. The RIBOSS core functions (in gold) incorporate previously published bioinformatic tools (in blue) in this pipeline. The analysis starts with mapping long reads using minimap2 (*14*). The RIBOSS transcriptome_assembly function takes long-read alignment files or a combination of long- and short-read alignment files as input (for StringTie) (*17*). ➀ The RIBOSS operon_finder and ORF_finder predict operons (for archaea and bacteria) and open reading frames (ORFs) from the top three frames of the newly assembled transcripts, ➁ align_reads maps ribosome footprints and paired RNA-seq data at the transcript isoform level (using STAR) (*18*), whereas ➂ quantify_transcripts estimate the abundance of each transcript isoform (using Salmon) (*19*), and riboprofiler assigns footprints by reading frames (using Ribomap) (*20*). Finally, the main RIBOSS function and statistical modules compare the translation of newly predicted ORFs with nearby annotated ORFs. Additional features include reporting the blastp hits and the Identical Protein Groups (*21, 22*) for the novel ORFs. The output files include publication quality figures, the annotation tracks for significantly translated ORFs, transcriptome assembly, and ribosome profiles adjusted to the P-site. These annotation tracks can be visualised using the UCSC Genome Browser, IGV, or Artemis (*23*–*25*).

The standard reference genome for *S* . Typhimurium is LT2. Long- and short-read transcriptome and ribosome profiling data mapped to this genome are available. We have used the transcriptome data from two independent studies. (i) A metatranscriptome data consisting of Nanopore long-read direct RNA sequencing (RNA-seq) and Illumina short-read RNA-seq data for a cocktail of *S* . *enterica* serovar Enteritidis, *Escherichia coli* O157:H7, and *Listeria monocytogenes* ( *12* ). These bacteria were grown in brain heart infusion media and romaine lettuce juice extract at 37°C for 24 h. (ii) An unpublished Nanopore long-read cDNA sequencing data for *S* . *enterica* ( *13* ). For ribosome profiling, the translatome data consists of ribosome profiling and matched short-read RNA-seq data for *S* . Typhimurium LT2 harvested at OD600 of 1, i.e. during the peak production of the SPI-1 transcriptional master regulator HilA ( *6* ).

We have mapped the long- and short-reads to *S* . Typhimurium LT2 reference sequences or concatenated sequences of *E. coli* O157:H7, and *L. monocytogenes* using minimap2 ( *14* ) and bowtie2 ( *15* ), respectively (Fig 1). The idea is to use the long- and short-read alignment files for *de no* transcriptome assembly. This is based on a typical gene annotation pipeline for model eukaryotes, where RNA-seq data from different sources have been used, including RNA-seq data from closely related species ( *16* ).

### RIBOSS predicts ORFs and operons in transcriptome assembly

The RIBOSS pipeline starts from the transcriptome_assembly function in the wrapper.py module. This function employs StringTie (*17*) to assemble transcriptomes using Nanopore or PacBio long-read alignment files, or a combination of long- and short-read alignment files. This function has presets for prokaryotic and eukaryotic transcriptome data. In particular, the preset for prokaryotes can suppress the assembly of spliced transcripts.

In this example, we have combined Nanopore direct RNA-seq and Illumina short-read RNA-seq data to assemble a metatranscriptome (*12*). We have also assembled a separate *Salmonella* transcriptome using a Nanopore cDNA sequencing data as mentioned above (*13*). We have merged these assembled transcripts with the LT2 reference gene annotation, generating a total of 4672 transcripts (Fig 2). The median length of the transcripts is 861 nt.

**Fig 2.**
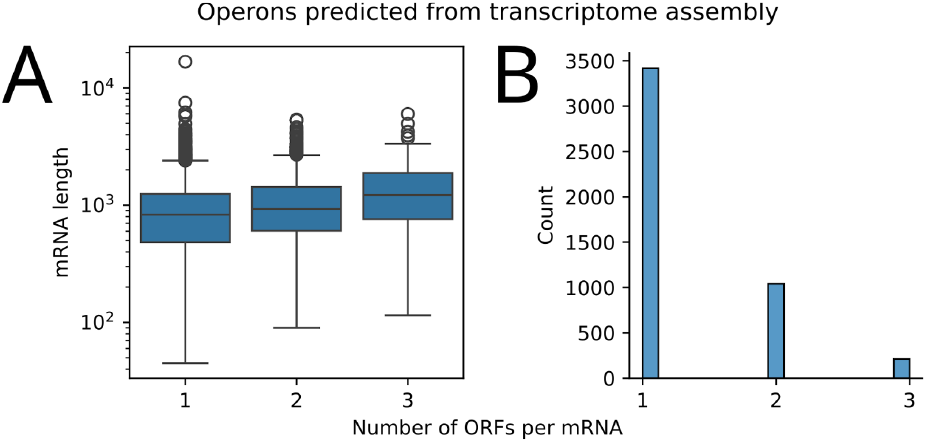
Operons detected through *de novo* transcriptome assembly. The RIBOSS transcriptome_assembly and operon_finder functions were used for this analysis. A total of 4672 transcripts were assembled by merging two independent transcriptome datasets with the reference gene annotation for *Salmonella enterica* serovar Typhimurium LT2. The datasets include a hybrid of Nanopore long-read direct RNA-seq and Illumina short-read RNA-seq metatranscriptome data (a cocktail of *S. enterica* serovar Enteritidis, *Escherichia coli* O157:H7, and *Listeria monocytogenes*) and Nanopore cDNA sequencing data (*S. enterica*). The median length of the assembled transcripts is 861 nt **(A)**, where 1254 of these transcripts harbour two or more annotated ORFs **(B)**.

We have used the RIBOSS operon_finder function to detect all possible ORFs in the top three frames of the assembled transcripts and matched them with annotated ORFs (Fig 1 ➀). By default, operon_finder (for archaea and bacteria) and ORF_finder (for eukaryotes) exclude annotated noncoding RNAs from the analysis. These two functions also remove the ORFs in-frame with annotated ORFs to avoid false positive results. As the ORFs found in a transcript isoform can overlap the annotated ORFs in a different isoform, the functions also remove them from the analysis. To achieve this, the functions convert the transcript level coordinates to the genomic level and compare them with the annotated ORFs.

We have found 1.3 genes per operon for *Salmonella* (Fig 2), which is similar to 1.4 genes per operon in *E. coli* (*26*–*28*). Notably, 1254 of these transcripts harbour two or more annotated ORFs, which is larger *E. coli* (788 polycistronic operons). This is expected as a previous study using PacBio long-read iso-seq has also extended the *E. coli* operon annotation (*29*).

Moreover, we have defined the ‘untranslated’ regions (e.g. intercistronic regions) and included these regions for ORF prediction. This approach can stimulate further investigation of these novel ORFs, e.g. their regulatory roles based on the operon model for RNA as a post-transcriptional circuit (*30*). This enables users to make the most of ribosome profiling to investigate translational control in polycistronic transcripts assembled using a hybrid of long- and short-reads (*26, 27*).

### RIBOSS compares the triplet periodicity for the ribosome footprints aligned to the transcriptome

The RIBOSS align_reads function maps ribosome footprints and the matched RNA-seq reads to the transcriptome using STAR (*18*) (Fig 1 ➁). Since the canonical footprint sizes are around 28 nt, the aligned footprint positions should be adjusted to the ribosomal P- or A-site (Fig 3A). Users can automate and customise this step using the analysed_footprints function. This function uses the alignment files to calculate the footprint positions relative to start and stop codons (Fig 3). This function also produces heatmaps for the aligned footprints by frame and the metagene plots for a chosen quality of footprint sizes.

**Fig 3.**
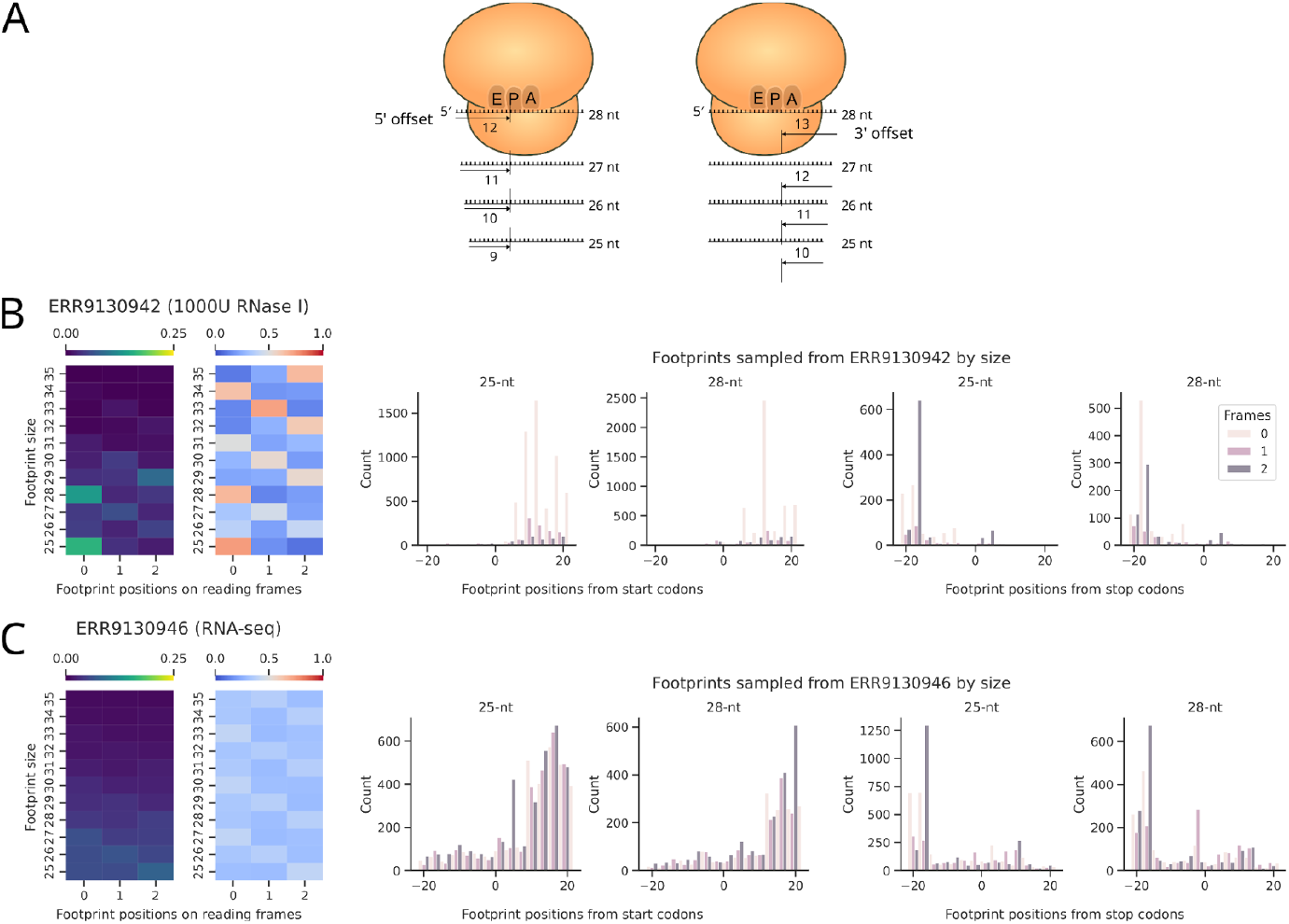
The RIBOSS analyse_footprints function evaluates ribosome footprints by size. **(A)** Users can adjust the aligned footprint positions to the P-site from the 5’ or 3’ end. Users can also customise the offset value for the footprint size of 28 nt, where this value will be automatically adjusted to offset different footprint sizes. **(B)** For the *Salmonella* data, the 5’ offset method produced consistent triplet periodicity for the footprint sizes 25 and 28 nt. **(C)** The matched RNA-seq reads lack triplet periodicity.

As the quality varies with footprint sizes, analysed_footprints in the footprints.py module uses chi-square post-hoc tests and/or odds ratio to evaluate the triplet periodicity of the footprints (similar to the section below). In this example, the high-quality footprints are 25-nt and 28-nt (Fig 3B). Such triplet periodicity is absent as expected in the RNA-seq data (Fig 3C). The aligned footprint positions were adjusted from the 5′ end, but users can adjust the footprints from the 3′ end (Fig 3A).

Through analysed_footprints, users can automate the selection of high-quality footprints to ensure that footprints can be correctly assigned by frame. The function also reports the offset values for the selected footprint sizes. This is critical for the riboprofiler function to correctly assign footprints to the coding and ‘noncoding’ regions using Ribomap (*20*). We have previously used Ribomap to analyse eukaryotic ribosome profiling data (*31*), as it can assign ribosome footprints to transcript isoforms.

### RIBOSS reports statistically significant novel translational events

The main RIBOSS function integrates upstream data and compares the translation of predicted ORFs and nearby annotated ORFs. However, the ribosome profiles along individual open reading frames are usually sparse except for highly expressed proteins. To handle data sparsity, the main function tallies the footprints aligned by frame for each ORF, creates 3×2 contingency tables for pairs of predicted and annotated ORFs (three frames × two ORFs), and compares the triplet periodicity using bootstrap chi-square tests (*32*). We have adopted a chi-square statistical module with bootstrap functionality (*33*) and developed a post-hoc test function based on a previous study (*34*). For a given pair of predicted and annotated ORFs, the main function calculates the odds of translation occurring for the predicted ORF using a 2×2 contingency table (footprints aligned to frames 1 and two other frames × two ORFs). The function uses stringent statistical thresholds to discover novel ORFs (odds ratio >1 and Bonferroni adjusted p-values >0.05 for bootstrap chi-square and post-hoc tests).

For the *Salmonella* data, 404 predicted ORFs (12 sORFs and 391 oORFs) passed the statistical thresholds (Fig 4). This represents ∼0.2% of all possible sORFs and oORFs that might be translated within the assembled transcripts. Of these predicted ORFs, one sORFs and 112 oORFs have significant blastp hits (Fig 4A and C). Our analysis rediscovered mgtP, i.e. the regulatory leader peptide of mgtCBR operon (*35*) (Fig 4E). Notably, mgtP has been annotated in a *Salmonella* genome (CP019649.1) but not in the LT2 reference strain. For oORFs, we found several interesting examples, including a partial plasmid entry exclusion gene traT overlaps the annotated traS (Fig 4G), and a partial O-antigen polymerase gene overlaps the annotated O-antigen polymerase rfbU gene (Fig 4H). These may be attributed to errors in genome assembly for LT2 or other strains. Among the predicted ORFs with no blastp hits, we found sORFs located upstream of the magnesium receptor mgtB and arginine deiminase arcA (Fig 4E and F).

**Fig 4.**
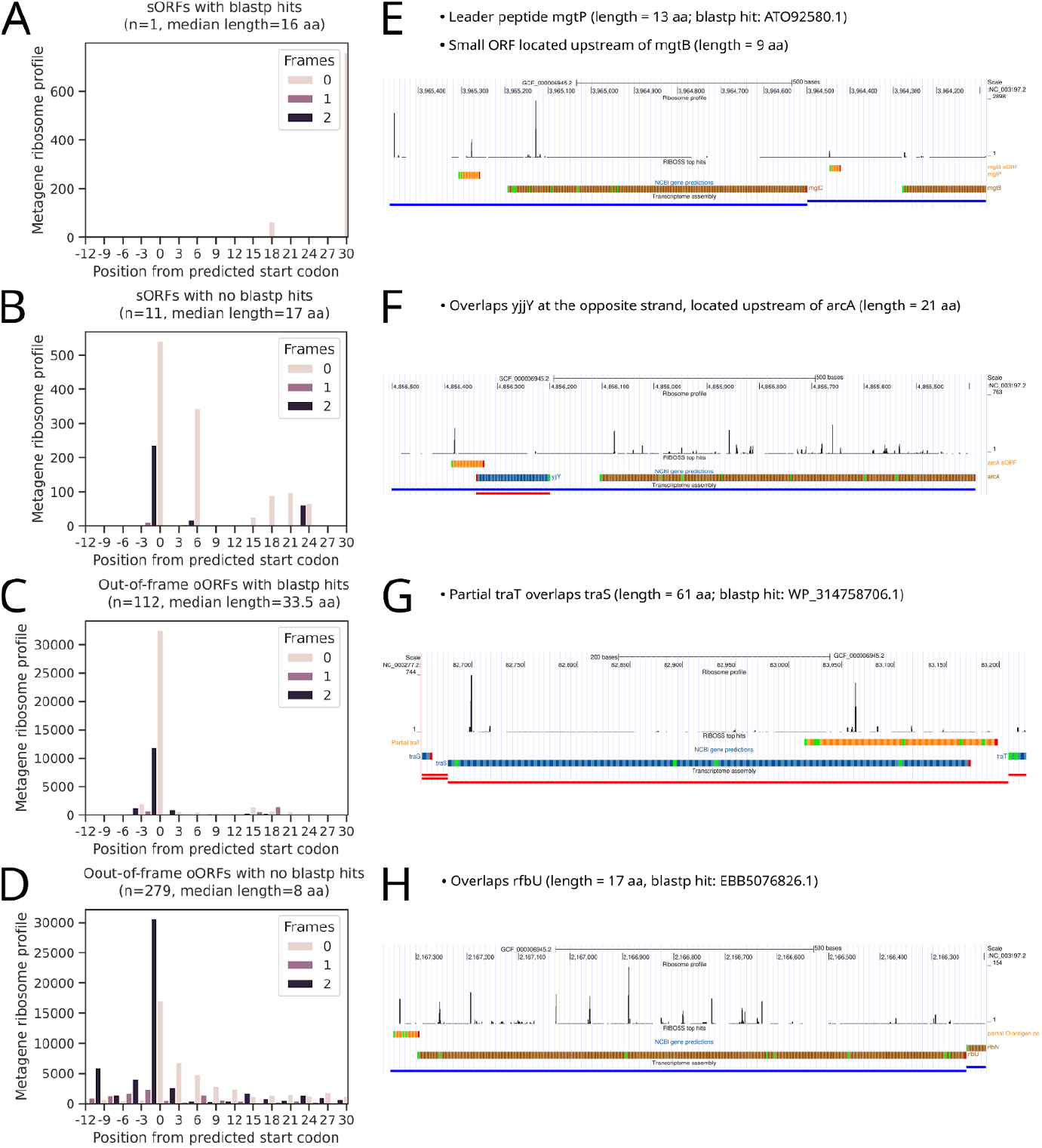
RIBOSS detects novel ORFs in the transcriptome assembled using a hybrid of long- and short-read data. RIBOSS compares the translation of predicted ORFs with annotated ORFs and reports statistically significant results together with blastp hits. These include sORFs (small ORFs) **(A, B)** and oORF (ORFs that overlap annotated ORFs out-of-frame) **(C, D)** detected in the *Salmonella* transcriptome. One of 12 sORFs and 112 of 391 oORFs have blastp hits, including the leader peptide mgtP **(E)**, a partial plasmid entry exclusion gene traT **(G)**, and a partial O-antigen polymerase **(H)**. mgtP is a regulatory leader peptide of mgtCBR operon. mgtP has been annotated in a *Salmonella* genome (CP019649.1) but not in the LT2 reference strain. No blastp hits were found for the sORFs located upstream of the magnesium transporter mgtB and arginine deiminase arcA **(E, F)**. The latter also overlaps yjjY at the opposite strand.

In summary, we have developed a Python package RIBOSS that enables the discovery of novel ORFs in transcriptome assemblies. Our new computational approach makes the best of long- and short-read RNA-seq technologies for high-quality *de novo* transcriptome assembly and ribosome profiling to reveal novel translational events.

## METHODS

### Data

Nanopore long-read direct RNA-seq and Illumina short-read RNA-seq data for a cocktail of *S. enterica* serovar Enteritidis (ATCC 13076), *E. coli* O157:H7 (ATCC 43895), and *L. monocytogenes* (ATCC 19115) were retrieved from PRJNA609733 (*12*). This metatranscriptome data corresponds to bacteria grown in brain heart infusion media and romaine lettuce (*Lactuca sativa* L. var. longifolia) juice extract at 37°C for 24 h. An independent Nanopore long-read cDNA sequencing data for *S. enterica* was retrieved from SRX20554650 (*13*).

The ribosome profiling and matched RNA-seq data for *S. enterica* serovar Typhimurium strain LT2 were retrieved from PRJEB51486 (*6*). This translatome data corresponds to LT2 cells harvested at OD600 of 1, i.e. during the peak production of the SPI-1 transcriptional master regulator HilA (*6*). The reference genome and GFF annotation files were downloaded from NCBI RefSeq (*36*).

### Sequence alignment

Long and short reads were aligned to *S*. Typhimurium LT2 reference sequences (chromosome NC_003197.2 and plasmid pSLT NC_003277.2) or concatenated sequences of LT2, *E. coli* O157:H7 (chromosome NZ_CP008957.1 and plasmid NZ_CP008958.1), and *L. monocytogenes* (NC_017544.1) using minimap v2.28 (*14*) and bowtie2 (*15*), respectively.

The ribosome profiling and matched RNA-seq reads were aligned to the newly assembled *Salmonella* transcriptome using build_start_index and align_reads employing STAR v2.7.11b (*18*). The build_start_index and align_reads functions are part of the wrapper.py module. SAMTools v1.21 were used to merge alignment files and convert the file formats (*37*).

### Transcriptome assembly

*De novo* transcriptome assembly was performed using a hybrid of Nanopore long-read and/or Illumina short-read alignment files. This was handled by transcriptome_assembly which employs StringTie v2.2.3 for transcriptome assembly (*17*), the UCSC Genome Browser’s KentUtils for file format conversion (*23*), and BEDTools v2.31.1 for transcript sequence extraction (*38*). The RIBOSS transcriptome_assembly function has a preset for prokaryotes to avoid spurious isoforms and spliced transcripts. The fraction of predicted transcript isoforms used is at least 0.1 of the most abundant transcript (default: 0.01). The minimum junction coverage used is 1000 for prokaryotes (to prevent prediction of spurious spliced transcripts) and 1 for eukaryotes (to include splicing) as default. The transcriptome_assembly function is part of the wrapper.py module.

### ORF and operon detection

The operon_finder and orf_finder functions detect ORFs in the top three frames of prokaryotic and eukaryotic transcripts, respectively. These include annotated ORFs and out-of-frame oORFs for prokaryotes and eukaryotes, sORFs for prokaryotes, and upstream and downstream ORFs for eukaryotes. To classify ORFs as such, PyRanges v0.1.2 was used to join the detected ORFs on genomic location (*39*). Unlike operon_finder, orf_finder was designed to handle the coordinates of splice junctions in eukaryotes. These functions are part of the orfs.py module.

### Ribosome footprint analysis

The triplet periodicity of ribosome footprints was examined using analyse_footprint. This includes downsampling the alignment files for ribosome profiling using pysam v0.22.1 (*40*), and adjusting footprints to P-site automatically (Fig 3A).

### Transcript quantification and ribosome footprint assignment

Transcript levels were determined by quantify_transcripts employing Salmon v1.10.3 (*19*). The ribosome footprints were assigned to reading frames and other ‘untranslated’ regions of transcripts using the C++ executable riboprof in Ribomap v1.2 (*20*).

### Sequence homology searching

The main RIBOSS function includes sequence homology search for significantly translated ORFs using the BLASTP module of Biopython (*22, 41*). The post-processing steps include parsing the blastp results using the topiary read.py module (*42*) and retrieving the Identical Protein Groups of the blastp hits using efetch in the NCBI Entrez module of Biopython (*21*).

### Data processing, statistical analysis, and visualisation

Pandas v2.2.3 and NumPy v1.26.4 were used for data processing (*43*–*45*). Plots were made using seaborn v0.13.2 and Matplotlib v3.9.2 (*46, 47*). Bootstrap chi-square tests were performed using a statistical module adopted from resamp v1.7.4 (*33*). Chi-square post-hoc tests were performed according to a previously published method (*34*) using SciPy v1.14.1 (*48*). Odds ratios were calculated using SciPy. The UCSC Genome Browser (*23*) was used to visualise the tracks generated by RIBOSS. The tracks include transcriptome assembly, ribosome profiles, and predicted ORFs in BED, bedGraph, and bigGenePred formats, respectively.

## DATA AVAILABILITY

RIBOSS is a Python package consisting of six modules. The source code is freely available on GitHub (https://github.com/lcscs12345/riboss). This package and its dependencies can be rapidly installed through the conda environment file using Miniforge3 v24.7.1-2 (*49*). The Jupyter notebook (*50*) and sequence alignment files to reproduce the results are available on Zenodo (https://doi.org/10.5281/zenodo.13997374). Raw sequencing datasets are available on NCBI BioProject/SRA (PRJNA609733 and SRX20554650) and ENA (PRJEB51486).

## ACKNOWLEDGMENTS

This work was funded by the Royal Society of New Zealand Te Apārangi, Marsden Fast-Start Grant (MFP-21-UOO-238), the University of Otago Research Grant (ORG 0122-0923, REDS 21954), and the University of Otago School of Biomedical Sciences Dean’s Bequest Fund (REDS 23704) to CSL.

## CONFLICT OF INTEREST

The authors declare no conflict of interest.

## Notes

https://github.com/lcscs12345/riboss

### Competing Interest Statement

The authors have declared no competing interest.

